# Differential encoding of odor and place in mouse piriform and entorhinal cortex

**DOI:** 10.1101/2023.10.05.561119

**Authors:** Wilson Mena, Keeley Baker, Alon Rubin, Shaun Kohli, Yun Yoo, Brice Bathellier, Yaniv Ziv, Shahab Rezaei-Mazinani, Alexander Fleischmann

**Affiliations:** Department of Neuroscience, Pasteur Institute, Paris, France; Department of Neuroscience and Carney Institute for Brain Science, Brown University, Providence, USA; Department of Neurobiology, Weizmann Institute of Science, Rehovot, Israel; Université Paris Cité, Institut Pasteur, AP-HP, Inserm, Fondation Pour l’Audition, Institut de l’Audition, IHU reConnect, F-75012 Paris, France; Center for Interdisciplinary Research in Biology (CIRB), Collège de France, Université PSL, CNRS, INSERM, 75005 Paris, France; Mines Saint-Étienne, Centre CMP, Département BEL, F-13541, Gardanne, France

**Keywords:** olfaction, odor encoding, spatial information, navigation, place encoding, pirifrom cortex, lateral entorhinal cortex

## Abstract

The integration of olfactory and spatial information is critical for guiding animal behavior. The lateral entorhinal cortex (LEC) is reciprocally interconnected with cortical areas for olfaction and the hippocampus and thus ideally positioned to encode odor-place associations. Here, we used mini-endoscopes to record neural activity in the mouse piriform cortex (PCx) and LEC. We show that in head-fixed mice, odor identity could be decoded from LEC ensembles, but less accurately than from PCx. In mice freely navigating a linear track, LEC ensemble activity at the odor ports was dominated by spatial information. Spatial position along the linear track could be decoded from LEC and PCx activity, however, PCx but not LEC exhibited strong behavior-driven modulation of positional information. Together, our data reveal that information about odor cues and spatial context is differentially encoded along the PCx-LEC axis.

**Significance statement:** For most animals, the sense of smell is essential for successfully navigating the environment to find food, shelter, and mates. However, how olfactory and spatial information is integrated in the brain to support olfactory-guided behaviors remains poorly understood. In mammals, candidate brain regions thought to support odor-place associations include the olfactory (piriform, PCx) cortex, entorhinal cortex, and hippocampus. We here characterize the activity of cells in the PCx and lateral entorhinal cortex (LEC) of freely moving mice in response to odor cues presented in a linear track. Using mini-endoscope microscopy and population coding analyses, we find that information about odors, spatial location, and behavior is differentially encoded along the PCx-LEC axis.

## Introduction

Odor cues in the environment serve as important spatial landmarks for animal navigation. In mammals, information about odors is represented in the olfactory cortex, while the hippocampus plays a key role in the encoding of spatial information ^1–8^. Olfactory cortex and hippocampus are strongly and reciprocally interconnected via the Lateral Entorhinal Cortex (LEC), and olfactory cortex, hippocampus and entorhinal cortex form the mammalian allocortex, suggesting a conserved link between neural representations of odor and space throughout evolution^9–11^.

Odors are detected by olfactory sensory neurons in the olfactory epithelium, and odor-evoked neural activity is initially transformed into spatio-temporal patterns of glomerular activity in the olfactory bulb^12,13^. Olfactory bulb mitral and tufted cells transmit odor-evoked neuronal activity to multiple cortical areas for olfaction, including the Piriform Cortex (PCx) and LEC^11,14,15^. The hippocampus has long been recognized for its role in encoding spatial and contextual information. Hippocampal place cells, for example, selectively fire when an animal traverses a particular location and are considered a neural substrate of spatial memory^4,6,16,17^.

Interestingly, recent findings have shown that odor cues at defined locations can serve as spatial landmarks and refine spatially-dependent neuronal activity in the hippocampus^18,19^. On the other hand, in rats performing an olfactory navigation task, PCx neurons robustly encode spatial information about the environment^20^. Together, these observations suggest that information about odors and their location in space is communicated across olfactory and hippocampal circuits.

The LEC is reciprocally interconnected with PCx and hippocampus and a prime candidate for integrating olfactory and spatial information. LEC receives direct olfactory inputs via projections from olfactory bulb mitral cells, and indirectly via PCx neurons^14,21,22^. The LEC receives spatial information via projections from hippocampus CA1 and subiculum, and indirectly via extensive reciprocal connections within the entorhinal-hippocampal network^9,23,24^. Therefore, a comparative analysis of odor-evoked and location-dependent neuronal activity along the PCx-LEC-hippocampal axis is critical for understanding neural circuit mechanisms for odor-place associations.

Here, we recorded neural activity in the anterior PCx and LEC, in mice passively exposed to odors, and in mice exploring odor cues in a linear track. We found that odor identity information was more accurately encoded in PCx than in LEC, independent of behavioral state. Place information, in contrast, was accurately represented in both LEC and in PCx. However, the representations of symmetric positions along the track were highly correlated in PCx, suggesting more schema-like representations of distances and actions. Together, our data suggest that information about odor and place are differentially encoded along the PCx-LEC axis.

## Results

### Odor representations in LEC and PCx in head-fixed mice

We stereotaxically injected Adeno-Associated Virus (AAV) expressing the calcium indicator GCaMP7^25^ into the LEC and anterior PCx of adult (6-8 week-old) mice. A Gradient Refractive Index (GRIN) lens was implanted above the injection site, and all behavioral and imaging experiments were performed between 6 to 12 weeks after lens implantation (**Supplementary Figure 1**). Visual inspection of individual imaging sites revealed well-identifiable cells exhibiting robust fluctuations in fluorescence (**Figure 1A-C**). Cell segmentation was performed using the Inscopix CNMFe algorithm.

**Figure 1.**
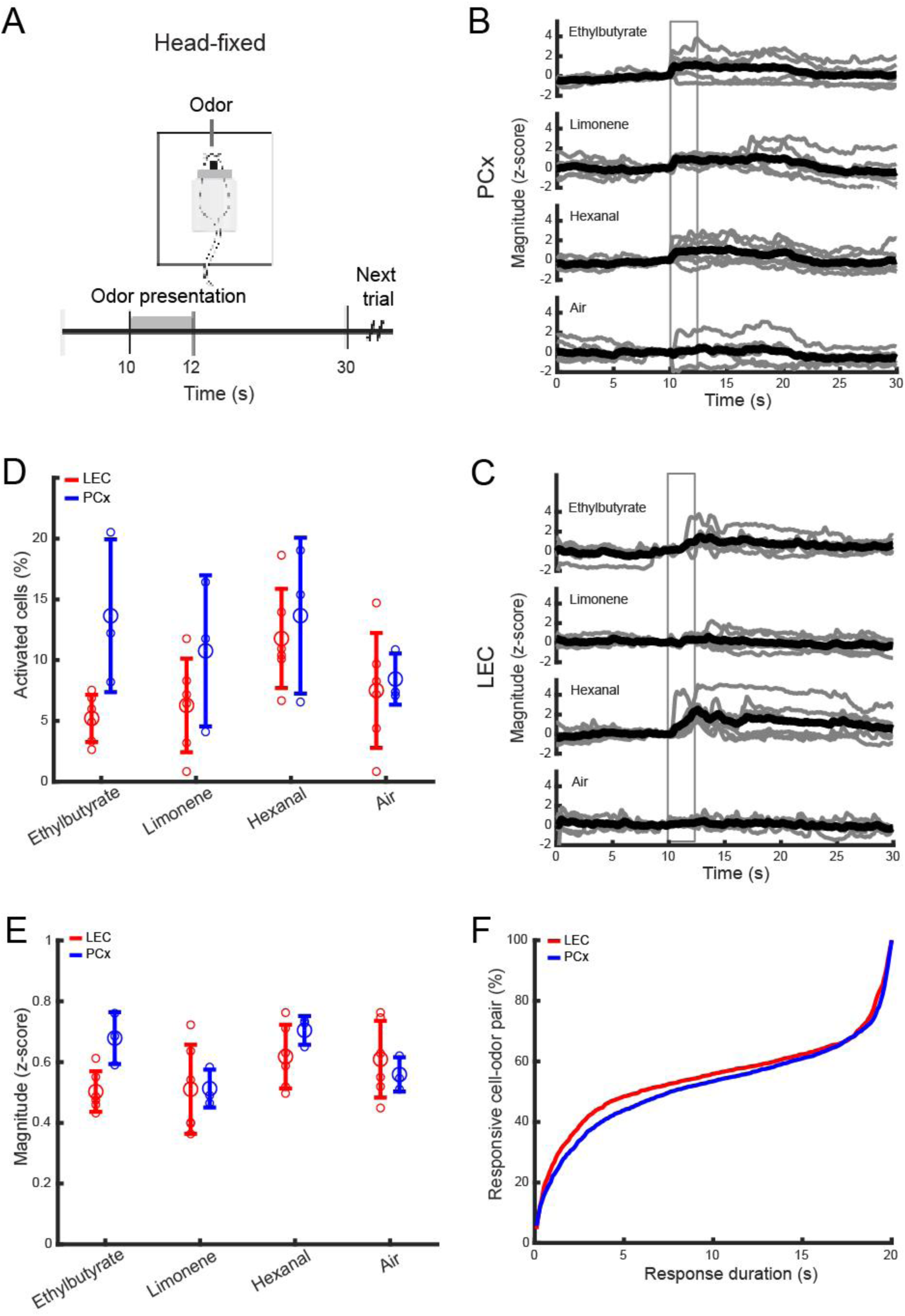
Calcium imaging of odor-evoked activity in LEC and PCx of head-fixed mice. **(A)** Head-fixed experimentation scheme and trial structure. **(B, C)** Example traces of 6 trials and their average, from one PCx and one LEC neuron, in response to three monomolecular odorants and clean air. Gray box: odor exposure window. **(D)** Percent of neurons activated in LEC (red) and PCx (blue). Large circles: average across 6 (LEC) and 3 (PCx) mice; small circles: data points from individual mice; bar: standard deviation. **(E)** Response magnitude in LEC and PCx, averaged across 6 (LEC) and 3 (PCx) mice. Large circles: average, small circles: data points from individual mice; bar: standard deviation. **(F)** Percent of responsive cell-odor pair as a function of response duration in LEC and PCx.

To compare basic odor response properties in LEC and PCx, we initially exposed head-fixed mice to ethylbutyrate, limonene, hexanal (0.3% vol./vol. in mineral oil), and clean air as a control (6 trials each, 30 sec inter-trial interval, see Methods; **Figure 1A-C**). We analyzed a total of 1235 LEC cells in 6 mice (mean ± SD: 205.8 ± 76.4), and a total of 611 PCx cells in 3 mice (mean ± SD: 203.6 ± 74.5).

We found that the percentage of odor-responsive cells in LEC was significantly lower than in PCx (LEC: mean ± SD: 7.69% ± 2.88, PCx: mean ± SD: 11.62% ± 2.52, Mann–Whitney *U* test; p = 0.03, **Figure 1D**). Furthermore, the response magnitude of cells in LEC appeared slightly lower than in PCx (LEC: mean ± SD: 0.55 ± 0.05 (z-score), PCx: mean ± SD: 0.61 ± 0.09 (z-score), p = 0.19, **Figure 1E**), and the distribution of odor-response duration (sec) was slightly shifted towards shorter responses in LEC than in PCx (Kolmogorov-Smirnov test; p = 0.25, **Figure 1F**). These data suggest that odor responses in LEC are less robust than in PCx.

We next asked whether LEC and PCx odor representations differ in how they encode odor identity information. We first compared the similarities of odor-evoked activity across trials and odorants. We calculated the pairwise cross-correlations between single-trial population response vectors during the 2 seconds of odor exposure, pooled across LEC and PCx imaging sites. Correlation matrices indicated that in LEC, overall trial correlations were lower than in PCx (**Figure 2A-D** and **Supplementary Figure 2**). Moreover, in contrast to PCx, repeated exposure to the same odorants did not elicit more correlated population activity than exposure to different odorants (LEC same odor correlation: mean ± SD: 0.11 ± 0.04, different odor correlation: mean ± SD: 0.09 ± 0.02, p = 0.18, PCx same odor correlation: mean ± SD: 0.44 ± 0.05, different odor correlation: mean ± SD: 0.34 ± 0.06, p = 5 × 10^-6^, **Figure 2C, D**).

**Figure 2.**
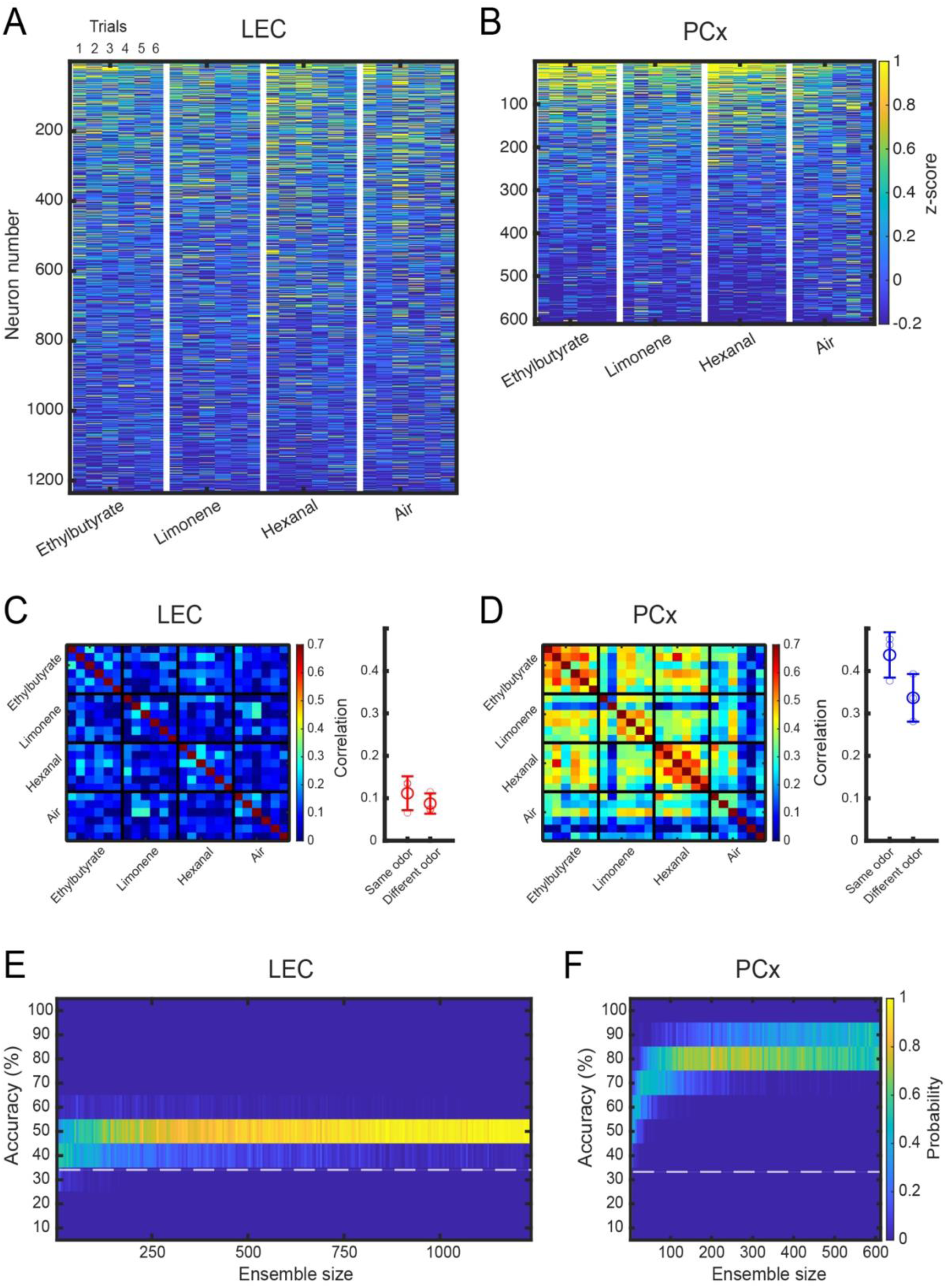
Representations of odor identity in LEC and PCx of head-fixed mice. **(A)** Heatmap of neural activity in LEC. **(B)** Heatmap of neural activity in PCx. **(C)** Left: Similarity matrix representing the pairwise correlation coefficients between neuronal activity population response vectors in LEC. Data obtained from 6 pooled mice (2 seconds odor-exposure window). Every small square represents a trial. A group of 6 trials constitutes an odor. Right: correlation of odor responses between repeat exposure to the same versus different odorants. Large circles: average, small circles: data points from individual mice; bar: standard deviation. **(D)** Left: Similarity matrix for neuronal activity population response vectors in PCx. Data obtained from 3 pooled mice (2 seconds odor-exposure window). Every small square represents a trial. A group of 6 trials constitutes an odor. Right: correlation of odor responses for same versus different odorants. Large circles: average, small circles: data points from individual mice; bar: standard deviation. **(E)** Accuracy of odor identity classification in pseudo-populations of increasing size in LEC. Distribution of the accuracy of odor classification using a linear SVM classifier trained on randomly sampled LEC ensembles of increasing size (2 seconds odor-exposure window). Total number of neurons: n = 1235. The white dotted-line shows the chance level. **(F)** Accuracy of odor identity classification in pseudo-populations of increasing size in PCx. Distribution of the accuracy of odor classification for PCx ensembles of increasing size (2 seconds odor-exposure window). Total number of neurons: n = 611. The white dotted-line shows the chance level.

To compare the accuracy of odor identity coding in LEC and PCx, we next used a linear Support Vector Machine (SVM) classifier (see Methods). Building classifiers with increasing numbers of randomly selected cells, we found that for similar ensemble size, average classification accuracy was significantly lower in LEC than in PCx (matched pseudo-population size of 610 neurons, LEC: mean ± SD: 43.5% ± 2.5, PCx: mean ± SD: 80% ± 2.9; p = 2.5 × 10^-34^, **Figure 2E, F** and **Supplementary Figure 2**). Furthermore, while accuracy in PCx reached peak levels within 0.55 seconds of odor onset, classification accuracy in LEC peaked after the 2 second window of odor exposure (**Supplementary Figure 2**).

### Odor representations in LEC and PCx in freely moving mice

We next asked how odor information was represented in LEC and PCx when freely moving mice sampled odorants at odor ports in a linear track. Mice were trained to sample an odorant at an odor port at the end of the track, where odor exposure was triggered when mice poked the odor port. Odor exposure lasted for two seconds, followed by a one-second delay before the water reward was delivered. Mice then moved to the other odor port to initiate the next trial (**Figure 3A**). Behavior and neuronal activity were recorded for a total of 48 trials: 12 trials of each odorant (ethylbutyrate, limonene, hexanal, clean air), 6 trials at each odor port (see Methods).

**Figure 3.**
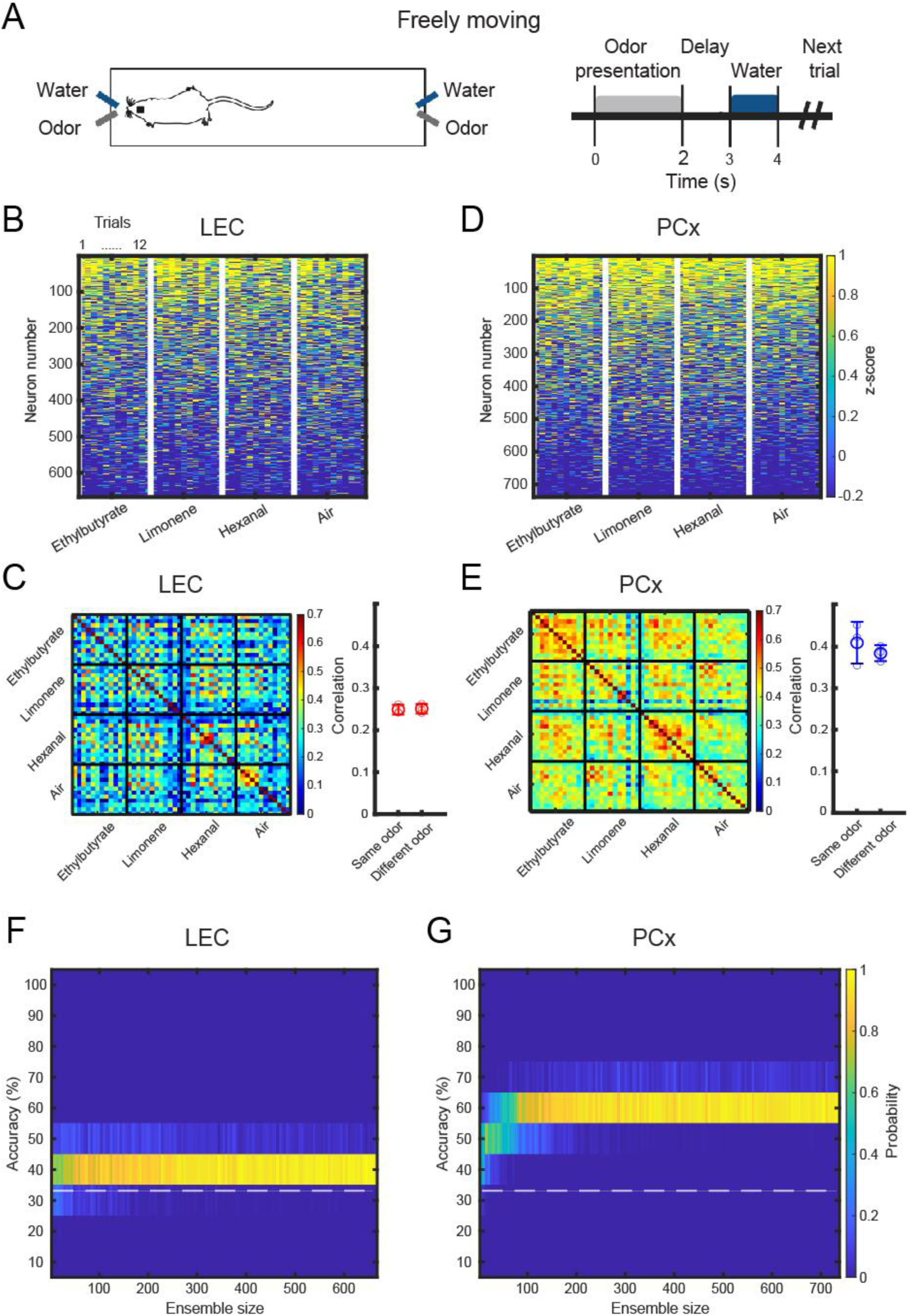
Representations of odor identity in LEC and PCx of freely moving mice. **(A)** Freely-moving experimentation scheme and trial structure. **(B)** Heatmap of neural activity in LEC. **(C)** Left: similarity matrix representing the pairwise correlation coefficients between neuronal activity population response vectors in LEC (2 seconds odor-exposure window). Data obtained from 3 pooled mice. Every small square represents a trial. A group of 12 trial constitutes an odor. Right: correlation of odor responses between repeat exposure to the same versus different odorants. Large circle: average, small circles: data points from individual mice; bar: standard deviation. **(D)** Heatmap of neural activity in PCx. **(E)** Left: similarity matrix for neuronal activity population response vectors in PCx (2 seconds odor-exposure window). Data obtained from 4 pooled mice. Every small square represents a trial. A group of 12 trial constitutes an odor. Right: correlation of odor responses for same versus different odorants. Large circle: average, small circles: data points from individual mice; bar: standard deviation. **(F)** Accuracy of odor identity classification in pseudo-populations of increasing size in LEC. Distribution of the accuracy of odor classification using a linear SVM classifier trained on randomly sampled LEC ensembles of increasing size (2 seconds odor-exposure window). Total number of neurons: n = 666. The white dotted-line shows the chance level. **(G)** Accuracy of odor identity classification in pseudo-populations of increasing size in PCx. Distribution of the accuracy of odor classification for PCx ensembles of increasing size (2 seconds odor-exposure window). Total number of neurons: n = 738. The white dotted-line shows the chance level.

We analyzed a total of 666 LEC cells in 3 mice (mean ± SD: 222 ± 91), and a total of 738 PCx cells in 4 mice (mean ± SD: 184 ± 118). Correlation matrices indicated that overall cross-trial correlations during the two second odor presentation in LEC increased in freely moving mice compared to head-fixed mice (**Figure 3B, C**). Furthermore, neural activity patterns in response to the same odorant were not more similar to each other than neural activity patterns in response to different odorants (same odor correlation: 0.24 ± 0.01, different odor correlation: 0.25 ± 0.01, p = 0.64, **Figure 3C**). PCx activity patterns similarly exhibited an overall increase in cross-trial correlations, independent from the odor stimulus (same odor correlation: 0.41 ± 0.05, different odor correlation: 0.38 ± 0.02, p = 3.2 x 10^-3^, **Figure 3D, E**). However, a linear SVM classifier accurately predicted odor identity in PCx, while classifier performance in LEC remained close to chance level (matched pseudo-population size of 665 neurons, LEC: mean ± SD: 36.8% ± 1.8, PCx: mean ± SD: 57.1% ± 1.8; p = 2.5 × 10^-34^, **Figure 3F, G** and **Supplementary Figure 3**). Together, these data suggest that odor identity coding in LEC is less accurate than in PCx, independent of whether mice are passively exposed to odors or actively sampling odors at odor ports in a linear track.

### Distinct representations of spatial location in LEC and PCx

The LEC receives strong inputs from the hippocampus and has been implicated in the integration of spatial and non-spatial information^9,26,27^. Furthermore, recent data show that in addition to odor information, spatial information is also encoded in the posterior PCx of rats^20^. We therefore next examined the representation of spatial information in LEC and PCx.

We reorganized cross-trial correlations in blocks of trials at each odor port, and we observed that in LEC, population activity was more highly correlated for trials at the same odor ports, independent of odor identity (same port correlation: 0.29 ± 0.03, different port correlation: 0.2 ± 0.03, p = 1.2 x 10^-7^). This odor port-specific correlation structure was not observed in PCx (same port correlation: 0.43 ± 0.1, different port correlation: 0.41 ± 0.1, p = 0.22, **Figure 4A, B**). Similar results were obtained for trials with clean air (LEC: same port: 0.29 ± 0.08, different port: 0.23 ± 0.1, p = 0.02; PCx: same port: 0.4 ± 0.12, different port: 0.37 ± 0.1, p = 0.12).

**Figure 4.**
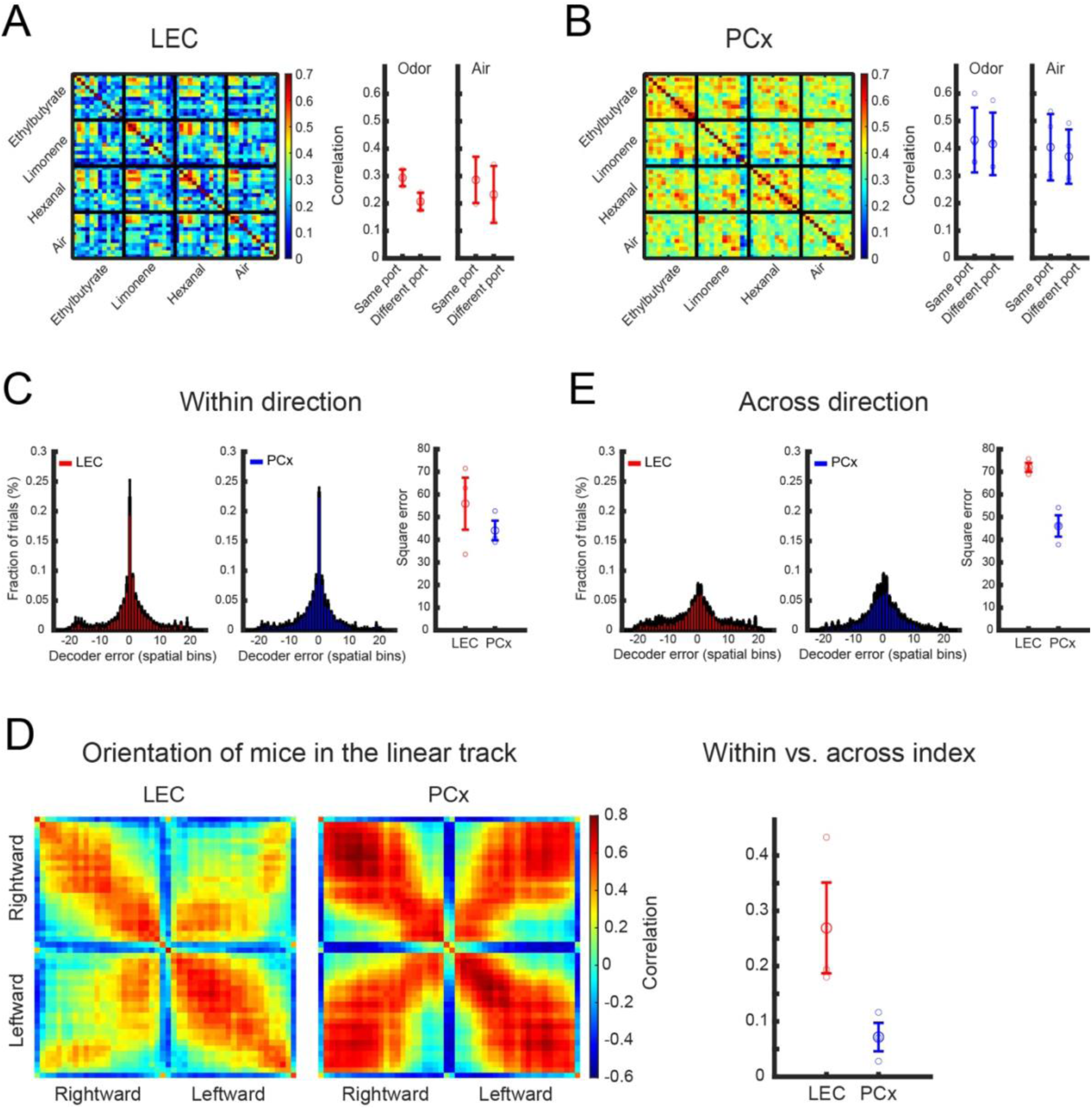
Representations of spatial location in LEC and PCx of freely moving mice. **(A)** Left: similarity matrix representing the pairwise correlation coefficients between neuronal activity population response vectors in LEC (2 seconds odor-exposure window), reorganized in trials at the left odor port (first 6 trials) and trials at the right odor port (second 6 trials). Right: Correlation of neuronal activity at the left odor port versus the right odor port, averaged across different odorants. The air shows the control. Large circle: average, small circles: data points from individual mice; bar: standard deviation. **(B)** Left: Similarity matrix for neuronal activity population response vectors in PCx (2 seconds odor-exposure window), reorganized in trials at the left and right odor port. Right: Correlation of neuronal activity at the left versus right odor port, averaged across different odorants. Large circle: average, small circles: data points from individual mice; bar: standard deviation. **(C)** Distributions of the decoder’s errors in inferring the position of mice along the linear track. The decoder was trained and tested on data from running in the same direction. **(D)** Pearson correlation between ensemble activity patterns across different spatial locations on the linear track, for data recorded in the LEC (left) and PCx (right). Ensemble activity patterns consist of concatenated epochs from a given location, separated according to the two running directions. **(E)** Distributions of the decoder’s errors in inferring the position of mice along the linear track. The decoder was trained on data from running in one direction and tested on data from running in the other direction.

We next tested the accuracy with which spatial position across the length of the linear track was represented in LEC and PCx. We defined a population vector for each position as the mean activity of each neuron at that position, and for each time frame, we identified the position for which the population vector had the highest Pearson correlation (see Methods). We found that decoders at single frame resolution demonstrate similarly high performance for both brain regions (**Figure 4C**). However, the analysis of the correlation structure of neuronal representations suggested differences in the encoding of spatial information relative to running direction. In LEC, the representation of a given position along the linear track was distinct for each running direction. In contrast, in PCx, the representation of symmetric positions along the linear track, relative to the odor ports, was highly similar for both running directions (**Figure 4D**). Consistent with this observation, we found that decoding of the trajectory phase was significantly more accurate in PCx than in LEC (differentiation index = (corr. same – corr. different) / (corr. same + corr. different); LEC: 0.27 ± 0.14; PCx: 0.07 ± 0.04, p (one-tailed t-test) = 0.04, **Figure 4E**.)

Together, these results reveal that spatial position along the linear track is accurately represented in LEC and PCx activity. However, LEC represents position largely independent from running direction, while PCx encodes a more schema-like representation of position and direction.

## Discussion

Memories of odors and places can be tightly intertwined, and odor-place associations are essential for animal navigation. The olfactory (piriform, PCx) cortex, lateral entorhinal cortex (LEC) and hippocampus form an evolutionarily conserved neural circuit that is ideally positioned to support odor-place associations^6,28^. We here recorded odor-evoked neural activity in PCx and LEC, using mini-endoscope calcium imaging. We compared odor representations in head-fixed mice passively exposed to odor, and in mice freely sampling odors at odor ports in a linear track. We found that odor identity was more accurately encoded in PCx than in LEC. Spatial information was accurately encoded in both LEC and in PCx. However, while LEC encoded spatial position continuously along the linear track, PCx preferentially represented behaviorally relevant positional information. Together, our data suggest that olfactory and spatial information is differentially processed within cortical olfactory circuits.

### Neural representations of odor identity are more robust in PCx than in LEC

In head-fixed mice, we found that odor-evoked neural activity was more robust in PCx than in LEC. The fraction of odor-responsive neurons and their response magnitudes were higher in PCx than in LEC, and responses were more prolonged. PCx neural activity patterns in response to re-exposure to the same odorant were more similar to each other than the neural activity patterns evoked by different odorants. In contrast, in LEC, odor-evoked neural activity was not correlated with odor identity. Finally, the accuracy of a linear SVM classifier to predict odor identity based on odor-evoked neural activity patterns was substantially higher in PCx than in LEC. Prior work has reported more accurate odor encoding in LEC than observed here^22,29,30^. The observed differences likely reflect differences in recording technology including temporal resolution, task design, and behavior. We here directly compared neural activity in PCx and LEC under the same experimental conditions. Thus, the relative differences in odor identity coding we observe likely reflect inherent differences between PCx and LEC.

Taken together, our results show that odor identity encoding was more robust in PCx than in LEC. This observation is consistent with well-established differences in neural circuit connectivity: PCx is the main target of olfactory bulb mitral and tufted cells. In contrast, LEC receives fewer direct olfactory bulb inputs, only from mitral cells^11,14,31^. Our data thus suggest that the robust reciprocal connections between PCx and LEC are likely to convey higher-order odor features, rather than information about odor identity alone.

### Task engagement does not improve odor coding in LEC

LEC is thought to integrate multimodal sensory and behavioral signals^6,9,23,32^, suggesting that LEC odor representations may be modulated by sensory and behavioral context. We therefore asked whether accurate odor identity representations in LEC require the active engagement of mice in a behavioral task.

We recorded odor-evoked activity while freely moving mice sampling odorants at odor ports in a linear track. We observed that odor-evoked neural activity at odor ports in the linear track was more highly correlated than under head-fixed conditions. However, correlated neural activity did not represent odor-specific information and was thus likely driven by behavioral and contextual signals at the odor ports. We found that the accuracy of odor identity coding in LEC was not increased with task engagement. Instead, odor identity coding decreased to chance levels in LEC, while it remained robustly above chance levels in PCx. Thus, sampling odors at odor ports in a linear track did not improve overall odor coding in LEC, relative to PCx. In our task, however, odor cues did not carry task-relevant information. In future experiments, it will be interesting to test whether tasks that rely on the specific association of odor with contextual spatial information engage LEC to represent odor information more accurately.

### Encoding of behaviorally relevant spatial information

Within the spatial and temporal limits of our experimental design, we found that the mouse’s position along the linear track could be decoded from LEC and PCx activity with similar accuracy. Our results extend the recent discovery of spatially tuned cells in the posterior PCx of rats^20^. Interestingly, however, our data suggest that LEC and PCx differ in how they represent behaviorally relevant spatial information.

We found that in LEC, the representation of a given position was distinct for each running direction, i.e. the representation of the same position differed for rightward and leftward running trajectories. In contrast, in PCx, the representation of symmetric positions along the linear track for opposite running directions was highly correlated, suggesting that PCx activity represented the position of the mouse relative to the start and end points of each track traversal, regardless of the running direction (i.e., the trajectory phase). This transformation of positional information into behaviorally relevant positional information is reminiscent of our recent findings in hippocampus and anterior cingulate cortex^33^. Determining the odor, spatial and behavioral tuning properties of individual PCx, LEC and hippocampal neurons will be critical for understanding the neural computations underlying odor-place associations. More generally, the ability to record from large ensembles of PCx, LEC and hippocampus neurons in freely moving mice provides a powerful approach for elucidating neural circuit mechanisms for the integration of sensory and contextual information.

## Materials and methods

### Mice

Adult C57BL/6J male mice (8 to 10 weeks old) were used in the study. All animals were grouped-housed with free access to food and water, in controlled temperature and humidity conditions, and exposed to conventional 12-hour light/dark cycles. All procedures in the experimental protocol were approved by the French Ethical Committee (authorization APAFIS#2016012909576100). Experiments were conducted in accordance with EU Directive 2010/63/EU. All efforts were made to reduce animal suffering and minimize the number of animals needed to obtain reliable results.

### Virus injection

All surgeries were performed under ketamine (80 mg/kg)-xylazine (1 mg/kg) anesthesia. The animal’s body temperature was maintained at 36°C using a feedback-based thermal blanket with a rectal probe (rodent Warmer X1, Stoelting). Mice were placed in a stereotaxic frame (David Kopf Instruments), and injections of 100 nl of GCaMP7s (pGP-AAV1-syn-jGCaMP7s-WPRE, Addgene plasmid #104487) diluted 1/3 in PBS were made at 50 nl/min at five locations separated by 200 - 250 µm. We used glass micropipettes and a manual injection system to deliver the virus, stereotaxic coordinates used for anterior PCx: A/P, +0.5 mm; M/L, -3.5 mm and D/V, -3.8 mm; for LEC: A/P, -3.15 mm; M/L, -4.15 mm and D/V, -2.5 mm. After surgery, mice were singly housed for one or two weeks without any manipulation.

### GRIN lens implantation

Two to three weeks after the viral injection, a GRIN lens of 0.5 mm diameter by 6.0 mm length (Inscopix, Palo Alto, CA) was implanted above the injection site to enable optical access to PCx or LEC neurons. First, a blunt needle was inserted and removed at 100 to 200 µm/min before the lens was implanted. The lens was sealed to the skull with dental cement (Super-Bond C&B, Sun Medical Co. Ltd.) and a low-viscosity composite (Flow-It ALC, Pentron). A head post, allowing transient head fixation of the animal to facilitate base plate positioning and miniature microscope mounting, was attached to the skull with Pi-Ku-Plast HP36 and low-viscosity composite (Flow-It ALC, Pentron). After lens implantation, animals were singly housed to prevent damage to the implant. No animals displayed evident motor or behavior abnormalities after the surgery. Two to three weeks following the surgery, a baseplate for the nVista microscope (v2.0, Inscopix, Palo Alto, CA) was mounted onto the animal’s head. In order to find a FOV with responsive cells, an odor puff was presented before baseplate implantation. After one to two weeks, mice started pre-training sessions.

Animals were habituated to the nVista miniature microscope attached to their head before the experimental session (pre-training phase 2; see below). GCaMP7s was excited with a light-emitting diode (LED) providing blue light at 475 mm, and fluorescence light was collected through a 535±25-nm emission filter and a complementary metal-oxide semiconductor (CMOS) camera chip, embedded in the miniature microscope (Inscopix, Palo Alto, CA). Following 15-20 min of free exploration of the linear track, recordings of neuronal activity were acquired at 20 Hz with a 50-ms exposure time, for 30-90 min.

### Behavior

During the entire behavioral task period, food was available *ad libitum* and animal weight was monitored daily. Water restriction (0.6-1 ml/day) was interleaved with a 12h *ad libitum* supply overnight every Friday. Mice performed behavior 2-3 days per week.

Pre-training: the linear track consisted of a Plexiglass structure 100 cm long, 8 cm wide, and 9 cm high. After 2 consecutive days of water restriction, mice were trained for 7-11 days to run back and forth along the track by giving them ∼10µl of water when nose poking on each end of the track. During pre-training, nose poking was followed immediately by odor presentation (1 sec) and the water reward (1 sec). Mice were kept under this regiment until they performed an average of 60 trials in 15 min on at least two consecutive days. After mice completed pre-training, a delay of 1 sec between odor presentation and water reward was added, and a 3D printed miniscope with similar dimensions and weight as the real one was implanted during training to habituate mice to run and nose poke with the miniscope attached.

### Calcium imaging and behavioral recordings

For experiments in the linear track, after pre-training, odorants (ethyl acetate, limonene, hexanal diluted to 0.3% vol./vol. in mineral oil) were introduced. The field of view (FOV), focus, signal gain, LED intensity, exposition time for GCaMP fluorescence were optimized for each mouse. FOVs selected during base plate implantation were the same for freely-moving recordings, and the same parameters for signal gain, LED intensity and exposition time were used. After setting up optimal imaging parameters, mice were put back into their home cage 20-40 minutes for miniscope habituation before the beginning of the task. Miniscope recordings and nose-poking detections were synchronized using the inscopix data acquisition box. A red LED light located next to the linear track was used to align the beginning and the end of calcium imaging and behavioral recordings.

For head-fixed recordings, mice were placed inside a modified Falcon tube and passively exposed to odor. Imaging parameters were adjusted to be the same between linear track and head-fixed regiments. The set of odorants, concentration and duration of odor presentation was the same as for freely-moving conditions. At the end of the recordings, the miniscope was removed and mice transferred to their home cage with free access to water and food. All brains were subsequently processed for histology.

### Histology

Histology was used to validate viral targeting of PCx and LEC. Mice were anesthetized with Euthasol (150 mg of pentobarbital/kg) and transcardially perfused with 10 ml of ice-cold phosphate-buffered saline (PBS) followed by 10 ml of 4% paraformaldehyde (PFA). Brains were dissected and post-fixed overnight in 4% PFA at 4°C and transferred to PBS. Coronal sections (200 µm) were prepared using a vibrating-blade Leica VT100S Vibratome. Sections were rinsed in PBS and incubated in PBS/0.1% Triton X-100 and Neurotrace counterstain (1:1000, ThermoFisher) at 4°C overnight, then mounted on SuperFrost Premium microscope slides (Fisher, cat# 12-544-7) in Fluorescent Vectashield Mounting Medium (Vector). Confirmation of both lens position and GCaMP expression was obtained using a Nikon A1R-HD confocal microscope. Overall, the success rate for this surgery, including methods development, was ∼80%. Most common failures were (i) scratched/chopped GRIN lens due to the animal removing the lens protection cap and (ii) excessive movement artifact or weak fluorescence signal on the FOV.

### Data structure

For all odor coding analyses, the output of Inscopix’s CNMF-based cell segmentation software was a 2-dimensional time series matrix, with cells in columns and calcium signals in rows. Head-fixed calcium signals consisted of 6 trials with 30 sec duration, for 4 stimuli, pseudo-randomly presented and with a 30 sec inter-trial interval. A 30 sec trial contained 10 sec pre-stimulus, 2 sec odor response, and 18 sec post-stimulus phases. For freely moving calcium signals, a trial contained a 1 sec pre-stimulus and 2 sec odor response phase. A total of 48 trials were organized into sets of 12 trials per odor, 6 trials at the right and 6 trials at the left odor port. For subsequent analyses, calcium traces were reorganized into a 4-dimensional dataset (cells, time, trials, odors).

### Preprocessing

We smoothened calcium signals using a sliding-window average of a length of 5 frames. Calcium signals had an arbitrary baseline. To normalize, we first calculated F_0_ by subtracting each signal from its mean value within an interval of 1 sec before stimulus presentation. We then z-scored signals by dividing F_0_ calcium signals by the standard deviation of all trials and odors.

### Cellular analysis

We calculated the percent of odor-responsive cells as the average of the fraction of active neurons across all trials for each odor. We selected activated neurons as those whose activity level was higher than the threshold of 3 times the standard deviation of the normalized baseline (1 sec before the stimulation onset), and remained above this threshold for at least 500 ms. To quantify response magnitudes, we averaged the magnitude of the z-scored signals of the activated cells, over the 2-second odor exposure period, across all trials of each odor, per mouse. Then, we calculated the average magnitude and standard deviation for all mice. Response duration was calculated as the time that a cell’s activity exceeded the threshold of 3 times the standard deviation of the normalized baseline, for a minimum of 15 consecutive frames. We averaged response durations over all the cells, trials and odorants.

### Population coding analysis

Pearson correlation: we quantified the similarity of PCx and LEC odor responses using Pearson correlation. We firstly calculated the average of calcium signals over the 2-second odor exposure period. We then concatenated all the trials of each odor per all cells. We computed Pearson correlation over all the cells across the ensemble of trials.

Odor classification: we quantified odor information encoded in PCx and LEC neural activity using a Support Vector Machine (SVM) classifier with a linear kernel. This classifier predicted odor identity based on response patterns of single trials. We used leave-one-out cross-validation, whereby every trial was excluded once from the computation of the kernel for assessment, while the rest of trials were used to train the classifier. Decoding accuracy was computed by classifying a given trial belonging to one of the stimulus groups consisting of 3 odors, with a chance level of 33.3%. The classifier was implemented using a 250 ms sliding window and classification accuracy was averaged across the 2-second odor exposure period.

Ensemble size classification: to determine classification accuracy as a function of ensemble size, we trained the classifier, as above, on odor responses from 5 randomly selected cells. We increased ensemble size by adding 5 more random cells, and tested odor classification. We continued this until the ensemble included all cells. This process was iterated 100 times, each time with different seed cells. To plot the distribution of classification accuracy per number of cells, we extracted the maximum classification accuracy from each cell group’s prediction curve within the 2 sec stimulus intervals. We plotted the distribution of maximum accuracy per cell group using histogram and scaled color image. We divided the distribution of cell groups by the number of iterations (100), to calculate the probability of maximum accuracy distribution per number of cells.

### Statistical analysis

All quantification and statistical analysis were performed using Matlab R2021 a. Statistical assessment was performed using non-parametric tests, as reported in the text. In all figures, error bars display standard deviation. 95% confidence interval was used in Supplementary Figure 2 for the error’s shaded area.

### Encoding of behaviorally relevant spatial information

For the linear track, we constructed a maximum correlation decoder to infer the position of the mouse at each frame based on the relations between the neuronal activity patterns during the exploration of the linear track; either in the same or the opposite running direction. For cross validation, frames on odd trials were decoded solely based on data from even trials and vice versa. We divided the 100 cm linear track into 24 bins, and we defined the activity pattern of each spatial bin as the vector of the activity rate per neuron at that bin. For each frame, we inferred the mouse’s position by maximizing the correlation of the activity pattern in that frame with the activity pattern of a bin (namely, the decoded position was the bin whose activity pattern had the highest correlation with that of the decoded frame). For this analysis, we considered only running epochs (velocity>2 cm/s and excluding the last 2 bins at each end of the tracks).

---

**Supplementary Figure 1.**
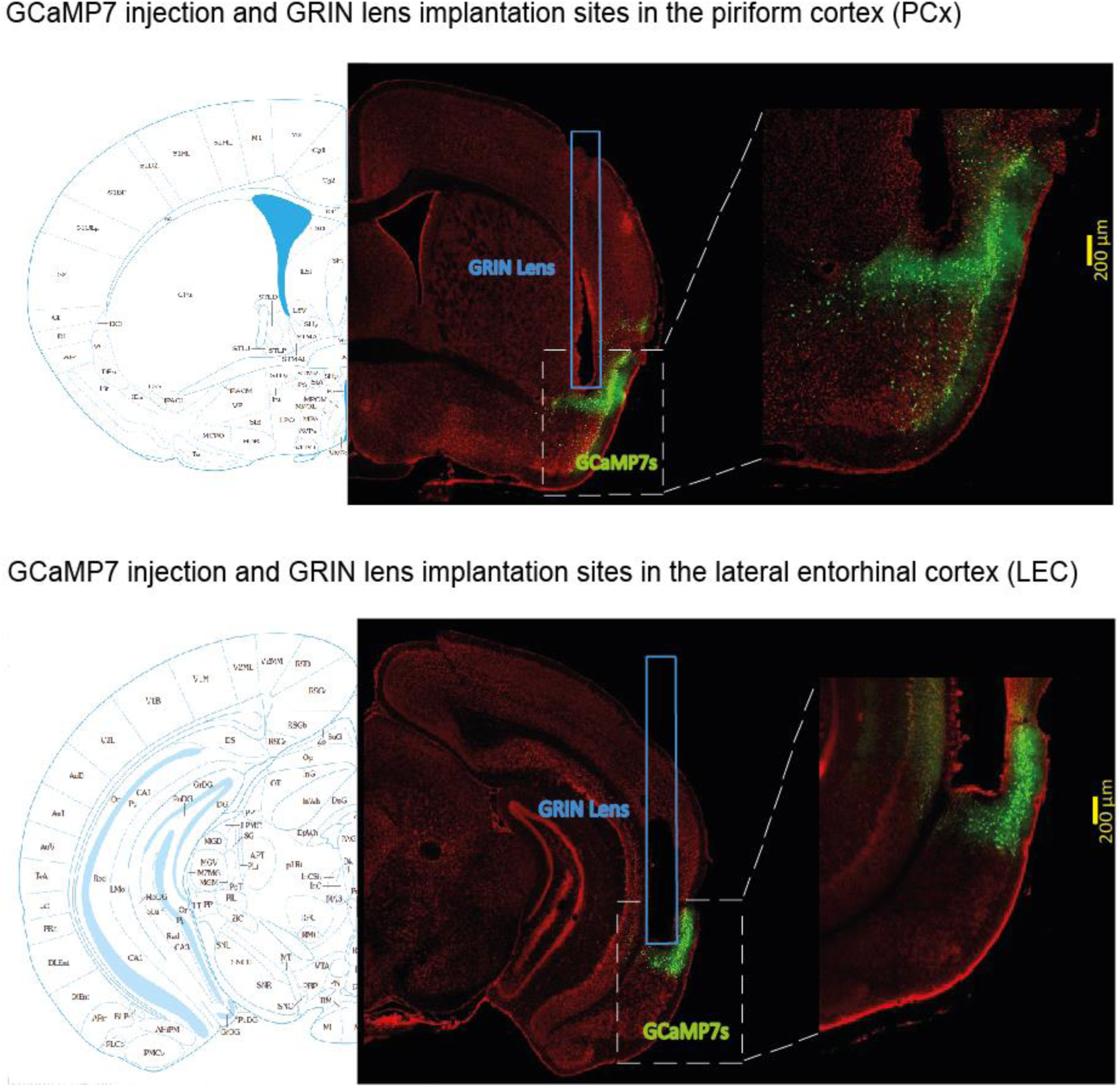
GCaMP7 injection and GRIN lens implantation sites in LEC and PCx. Top left: Atlas schematics of coordinates of viral injection and GRIN lens implantation for the piriform cortex. Top middle: Representative GCaMP7s injection and GRIN lens implantation sites in the piriform cortex. Top right: An enlarged view of the GCaMP7s injection and GRIN lens sites in the piriform cortex. Scale bar: 200 µm represents the approximate imaging distance below the GRIN lens. Bottom left: Atlas schematics of coordinates of viral injection and GRIN lens implantation of the LEC. Bottom middle: Representative GCaMP7s injection and GRIN lens implantation sites in the LEC. Bottom right: An enlarged view of the GCaMP7s injection and GRIN lens sites in LEC. Scale bar: 200 µm represents the approximate imaging distance below the GRIN lens.

**Supplementary Figure 2.**
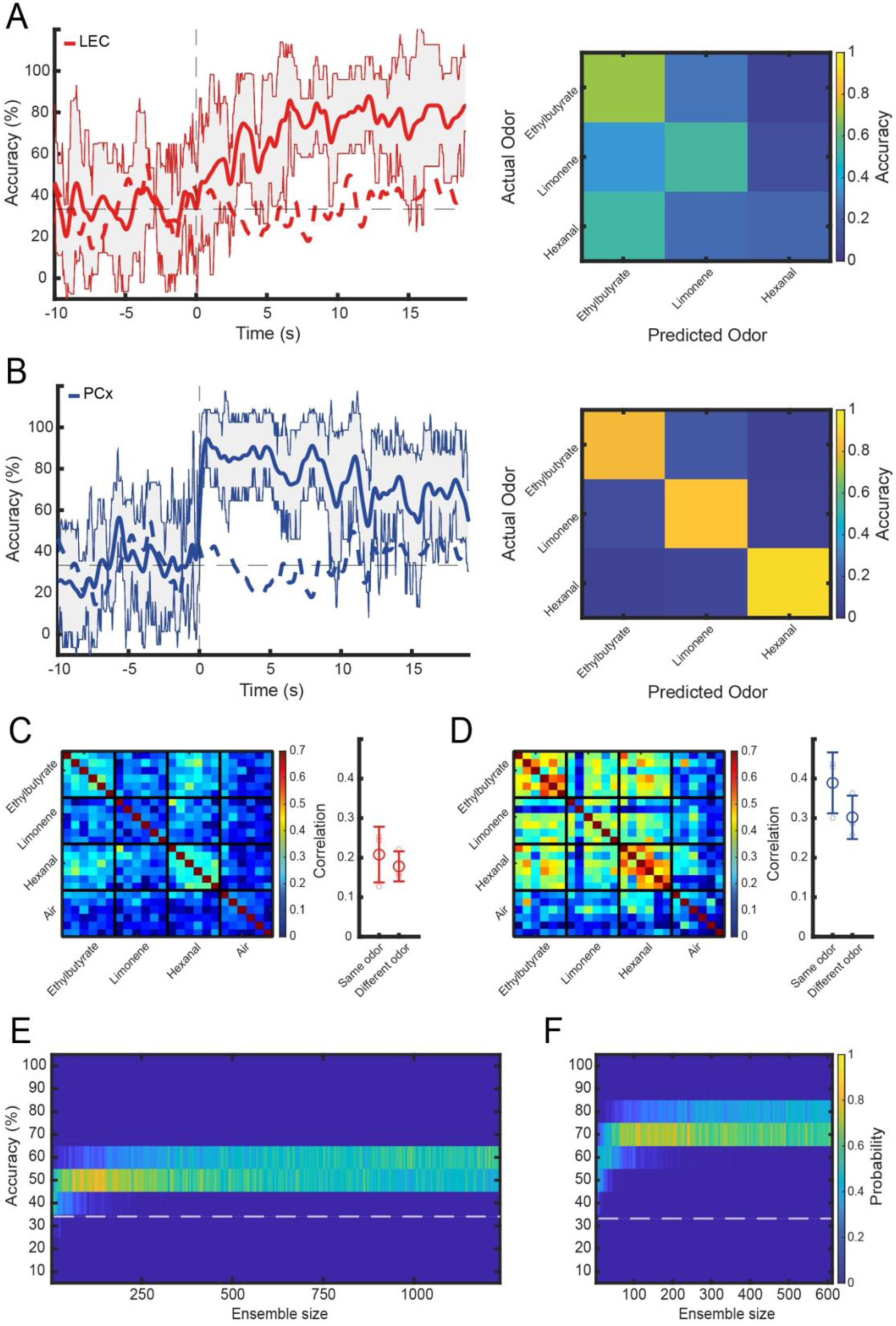
Dynamics of odor identity encoding in LEC and PCx in head fixed. **(A,B)** Left: Accuracy of odor identity classification in LEC (A) and PCx (B) over time, for head-fixed mice. Time 0 indicates odor valve opening. Odor encoding is less accurate and delayed in LEC compared to PCx. Shaded area indicates 95% confidence intervals for the mean. Right: Confusion matrix summarizing the performance of the SVM classifier trained to discriminate the odorants. Decoding accuracy for each odor, averaged over a 5 second time window. **(C)** Left: Similarity matrix representing the pairwise correlation coefficients between neuronal activity population response vectors in LEC. Data obtained from 6 pooled mice (5 seconds odor-exposure window). Every small square represents a trial. A group of 6 trials constitutes an odor. Right: correlation of odor responses between repeat exposure to the same versus different odorants. Large circles: average, small circles: data points from individual mice; bar: standard deviation. **(D)** Left: Similarity matrix for neuronal activity population response vectors in PCx. Data obtained from 3 pooled mice (5 seconds odor-exposure window). Every small square represents a trial. A group of 6 trials constitutes an odor. Right: correlation of odor responses for same versus different odorants. Large circles: average, small circles: data points from individual mice; bar: standard deviation. **(E)** Accuracy of odor identity classification in pseudo-populations of increasing size in LEC. Distribution of the accuracy of odor classification using a linear SVM classifier trained on randomly sampled LEC ensembles of increasing size (5 seconds odor-exposure window). Total number of neurons: n = 1235. The white dotted-line shows the chance level. **(F)** Accuracy of odor identity classification in pseudo-populations of increasing size in PCx. Distribution of the accuracy of odor classification for PCx ensembles of increasing size (5 seconds odor-exposure window). Total number of neurons: n = 611. The white dotted-line shows the chance level.

**Supplementary Figure 3.**
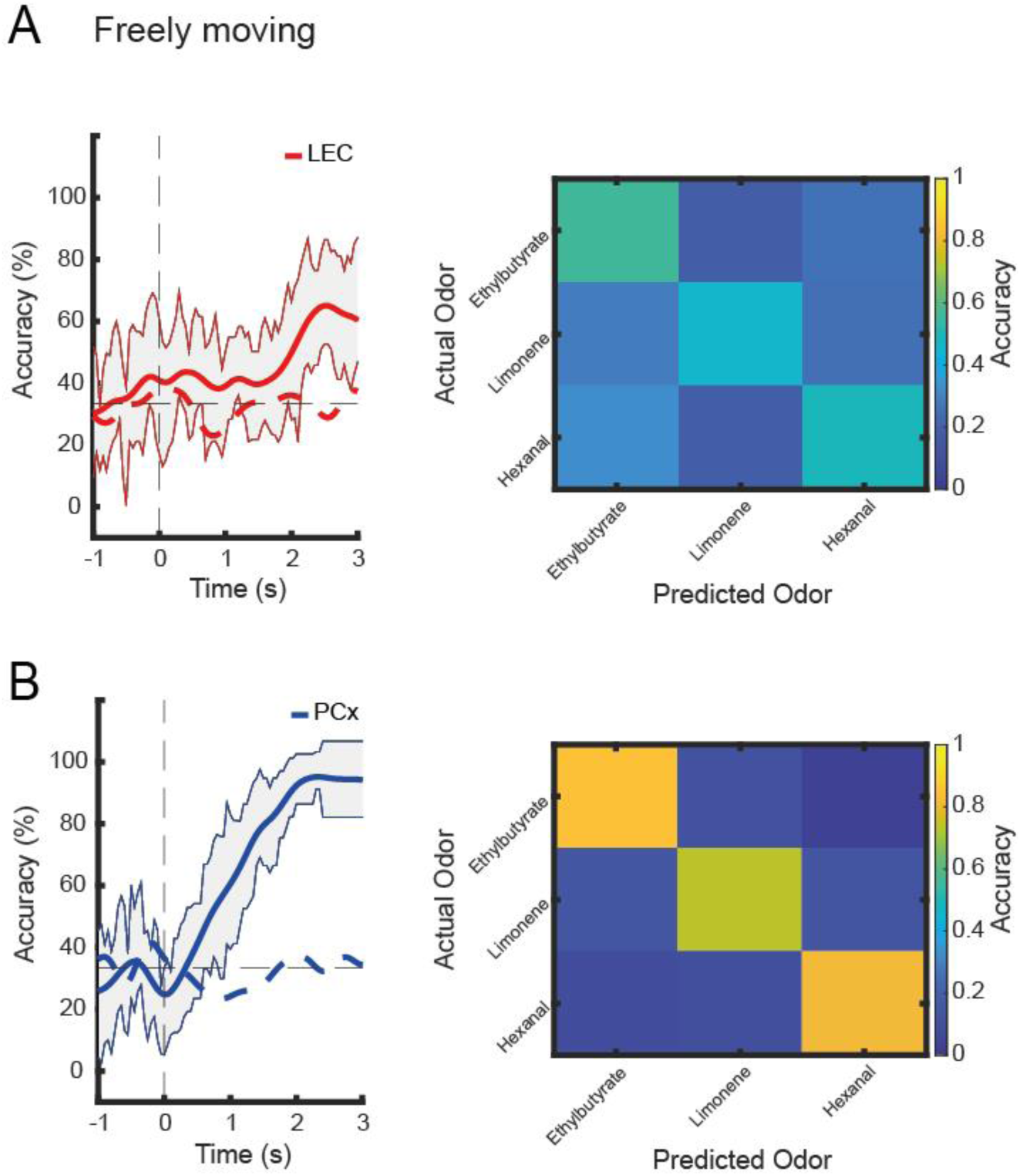
Dynamics of odor identity encoding in LEC and PCx in freely moving. **(A, B)** Left: Accuracy of odor identity classification in LEC (A) and PCx (B) over time, for freely-moving mice. Time 0 indicates odor valve opening. Odor encoding is less accurate and delayed in LEC compared to PCx. Shaded area indicates 95% confidence intervals for the mean. Right: Confusion matrix summarizing the performance of the SVM classifier trained to discriminate the odorants. Decoding accuracy for each odor, averaged over a 5 second time window.

## Acknowledgements

We thank members of the Fleischmann lab for critical comments on the manuscript. Work in the AF lab was supported by grants from the NIH (NIDCD R01DC017437), and the Robert J and Nancy D Carney Institute for Brain Science, Carney Institute computational resources used in this work were supported by the NIH Office of the Director grant S10OD025181. The authors thank Cindy Poo for valuable feedback, Michaël Zugaro and Karim Benchenane for their help with experiments, and Tuan Pham for help with data analysis and visualization.

## Author contributions

WM, SRM, and AF conceptualized the study with input from all authors. WM conducted all experiments and initial imaging data processing. KB and YY performed histological analyses and additional imaging data processing. SRM, BB and AF performed odor coding analyses, SK, AR and YZ performed place coding analyses. SRM and AF wrote the manuscript with input from all authors.

## Competing interests

Authors declare that they have no competing interests.

## Notes

### Competing Interest Statement

The authors have declared no competing interest.

### Summary of Updates

Text corrections and additional supplementary data.

